# Cell-specific occupancy dynamics between the pioneer-like factor Opa/ZIC and Ocelliles/OTX regulate early head development in embryos

**DOI:** 10.1101/2022.12.15.519123

**Authors:** Kelli D. Fenelon, Fan Gao, Priyanshi Borad, Shiva Abbasi, Lior Pachter, Theodora Koromila

## Abstract

During development, embryonic patterning systems direct a set of initially uncommitted pluripotent cells to differentiate into a variety of cell types and tissues. A core network of transcription factors, such as Zelda/POU5F1, Odd-paired (Opa)/ZIC and Ocelliless (Oc)/OTX, are conserved across animals. While Opa is essential for a second wave of zygotic activation after Zelda, it is unclear whether Opa drives head cell specification past gastrulation onset, in the *Drosophila* embryo. Our hypothesis is that Opa and Oc are interacting with distinct cis-regulatory regions for shaping cell fates in the embryonic head. Using super-resolution microscopy and epigenomic meta-analysis of single cell RNAseq datasets we find that *opa*’s and *oc*’s overlapping expression domains are dynamic in the head region, with both factors being simultaneously transcribed at the blastula stage. However, analysis of single-embryo RNAseq data reveals a subgroup of Opa-bound genes to be Opa-independent in the cellularized embryo. Interrogation of these genes against Oc ChIPseq combined with in situ data, suggests that Opa is competing with Oc for the regulation of a subgroup of genes later in gastrulation. Specifically, we find that Oc binds to late, head-specific enhancers independently and activates them in a head-specific wave of zygotic transcription, suggesting distinct roles for Oc in the blastula and gastrula stages.

## Introduction

Early embryos undergo multiple waves of zygotic genome activation regulated by well-orchestrated transcription factor (TF) networks that lead to organogenesis^1–3^. The roles of *Drosophila* transcriptional activators such as Bicoid (Bcd, PITX2 human ortholog)^4^, Zelda (Zld, POU5F1 human ortholog)^5^, Odd-paired (Opa, zinc finger protein of the cerebellum 3 (ZIC3), human ortholog)^6^, and Ocelliless (Oc, OTX2 human ortholog)^7^, are largely conserved across animals^8^. The critical nodes of the regulatory networks are promoter regions which are required for gene transcription; however, a significant part of transcriptional regulation occurs via the action of distal cis-regulatory modules, enhancers, where TFs bind to activate or repress target genes^9–11^. Furthermore, a gene with a complex expression pattern may have multiple enhancers active at any particular stage, each responsible for a discrete spatiotemporal aspect of the gene’s expression. Most enhancers can act in sequence to support gene expression at different developmental points^9,12,13^.

Within the first two hours of embryo development, transcriptional regulation shifts from maternally loaded control to zygotic regulation^5,14^. Specifically, there is a hand-off from Zld to Opa early in zygotic genome activation^3,15^. Additionally, Bcd can bind to inaccessible chromatin on its own at high concentrations anteriorly^16–18^, but requires Zld to do so at low concentrations in the posterior^19^. Later in embryogenesis, more confined, broadly expressed transcription factors, like Opa, drive the transcriptional landscape to undergo a dramatic shift to prepare the syncytial nuclei for cellular sovereignty rounding out the blastula stage and transitioning the embryo into gastrulation^20^. Opa and Oc begin their expression prior to gastrulation with known head and thorax developmental functions^21,22^. A previous study showed that a group of Anterior-Posterior axis (AP) enhancers are initially activated by Bcd, and later activation is transferred to Oc via a feed-forward relay that involves sequential binding of the two proteins to the same DNA motif^23^. However, the differential action between the pioneer factor Opa and Oc on epigenetic timing and levels of gene expression in the embryonic head is still unknown.

A vast and growing number of genomics and transcriptomics studies have produced a panoply of ChIPseq, whole embryo and single-cell RNAseq^24^, and other genomics datasets available to the public. Our novel data were compared to these public datasets to reveal mechanisms of transcriptional control otherwise undetectable. We found that balance between pioneer factors (Opa) and localized activators (Oc) is important in regulating timing of gene expression pre-cellularization. Further, analysis of scRNAseq data reveals *opa/oc* coexpressing cells at stage 5 (St5, pre-and during cellularization), but not later, and enrichment of several known neural developmental genes in cells containing both *opa* and *oc* transcripts. Interrogation of these genes against Opa, Oc, Bcd, and Zld ChIPseq datasets, expression databases, and published enhancer data suggests a high likelihood that *oc* and/or *opa* regulate their expression at distinct timepoints during development.

Transcription factors which share homology and function between *Drosophila* and mammals present optimal utility in studying developmental phenomena with both broad and specific impacts, e.g. very early brain development and the impact of those cell differentiation pathways and environmental factors on complex neurological disorders like autism spectrum disorder (ASD)^25^. Gene replacement experiments show that the *Drosophila oc* gene and orthologous mammalian *Otx2* gene are functionally equivalent^7,26^. In head development, different levels of OTX protein are required for the formation of specific subdomains of the adult head^21,26^. Also, ZIC2, has been known to play major roles in neural progenitors regulation^27,28^. Our *in situ* hybridization data show that both Opa and Oc affect gene expression dynamics at different timepoints during early embryo development. Thus the *Drosophila* embryo is a powerful system to study gene regulatory mechanisms involved in head development.

## Results

### Opa and Oc co-occupy genomic loci and embryonic region pre-gastrulation

We first sought to investigate the expression dynamics of *opa* and *oc* at 4 timepoints, just before cellularization (Stage 5 early: nc14B), as cellularization occurs (Stage 5 late: nc14C and nc14D) and at the onset of gastrulation (Stage 6). At stage 5 early (St5E), we found that both *oc* (anterior, future head, region of the embryo) and *opa* (broad central region of the embryo) are expressed as previously published (Figure 1A). Further investigation revealed that upon commencement of their transcription, *oc’s* and *opa’s* expression domains overlap in the posterior portion of the future head region (Figure 1A). This overlapping domain remains through cellularization but shrinks as cellularization ends and gastrulation begins (Figure 1A,B, Supplemental Figure 1A,B,C)^30^. Quantitative analysis of normalized fluorescent signal reveals an apparent posterior shift of the anterior boundaries of both *oc* (4% to 8%) and *opa* (16% to 25%) expression domains with the overlapping region shrinking from around 16% to 7% of the AP body axis between St5E and initiation of gastrulation (Figure 1A,B). To investigate whether Opa and Oc may cooperate to affect gene expression in this overlapping region, we interrogated publicly available ChIPseq data for Opa and Oc genomic binding^3,23^. Intriguingly, we found that a large majority of Oc ChIPseq peaks overlap with Opa peaks with 85% of Oc peaks during early cellularization and 83% of Oc peaks at the onset of gastrulation coinciding with Opa peaks (Figure 1C).

**Figure 1.**
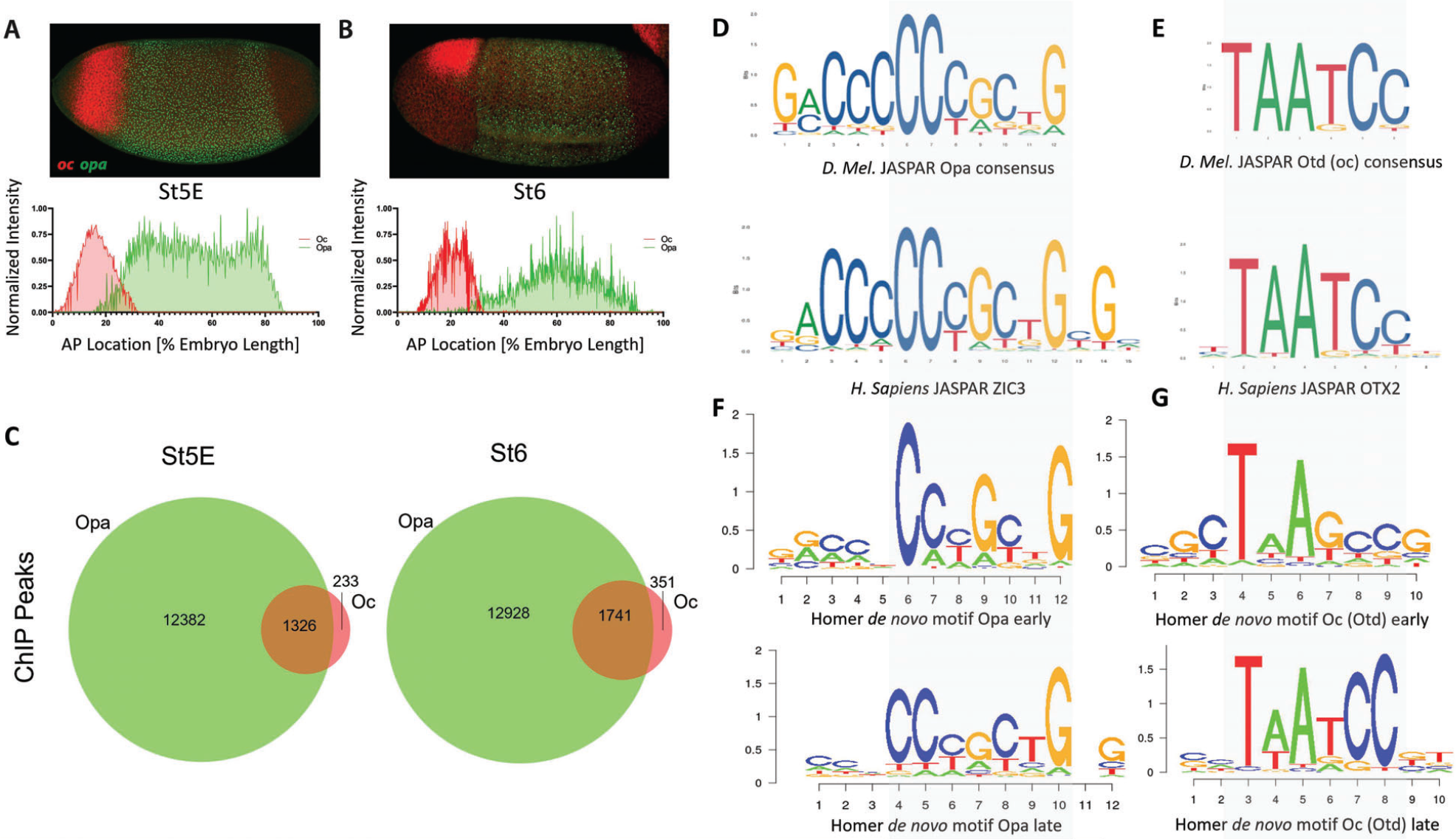
*opa* and *oc* dynamic overlap within the developing embryo. (A) At nc14B, a clear overlap between *oc* and *opa* domains can be observed (above) as is visualized graphically by batch plotting of AP FISH image fluorescence intensity below (n=4,3). (B) By initiation of gastrulation, *oc* and *opa* expression domains become nearly distinct, as verified by batch plotting of AP FISH image fluorescence intensity (n=7,7). (C) Venn diagram representing ChIP peaks for Opa (green) and Oc (red) at early and late stages. (D-E) Consensus binding data from JASPAR show that the *Drosophila* Opa (D), and Oc (E) consensus binding sites are conserved in Human. (F-G) Homer *de novo* motifs for Opa (F) and Oc (G) for early and late stages.

The existence of a shrinking overlap in expression of *opa*/*oc* implies existence of a narrow spatiotemporal window where these transcription factors may be capable of simultaneous occupation of enhancer regions in a small group of early embryonic cells to initiate a unique, nascent lineage, whilst maternal transcription factors, e.g. Bicoid (Bcd) and Zelda (Zld), phase out through cellularization in favor of zygotically expressed transcription factors (Supplemental Figure 1D). We next performed *de novo* motif analysis on these datasets to confirm consensus preservation between these datasets and the published, evolutionarily conserved JASPAR^31^ motifs for Opa/ZIC3 and Otd/OTX2 (Figure 1D-G). Interestingly, *de novo* motif analysis for Opa and Oc (Figure F,G) revealed cleaner, more precise motifs for St5L than St5E suggesting potential binding site competition early.

### Opa and Oc binding resolves post-cellularization

As we hypothesized distinct and cooperative roles for Opa and Oc transcription factors during cellularization, we next wished to investigate the binding dynamics of Opa and Oc during MBT. Toward this end, we further interrogated our HOMER *de novo* motif analyses from St5E and St6 embryos.

Motifs from *de novo* analysis of Oc and Opa non-overlapping peaks revealed the characteristic motifs for each transcription factor, TAATCC and CCCGCTG, respectively in both early and late datasets (Figure 2A,B,D,E). As expected, when the ratio of these transcription factors to maternal factors and total peak counts are low, the early dataset did not produce meaningful aggregations of predicted motifs around Oc peak loci (Supplemental Figure 2.1B). However, at the onset of gastrulation, there are significant peaks to reveal aggregation of Oc and Opa motifs at sites of their respective ChIPseq peaks (Figure 2C,F). Motifs from *de novo* analysis of Oc and Opa overlapping peaks also included the characteristic motifs for each transcription factor in both early and late datasets (Figure 2G,H). Interestingly, there appears to exist a shift in Oc and Opa motif proximity to their individual ChIPseq peaks from clustering more around Opa peaks early to clustering more around Oc peaks at the later stage (Figure 2I and Supplemental Figure 2).

**Figure 2.**
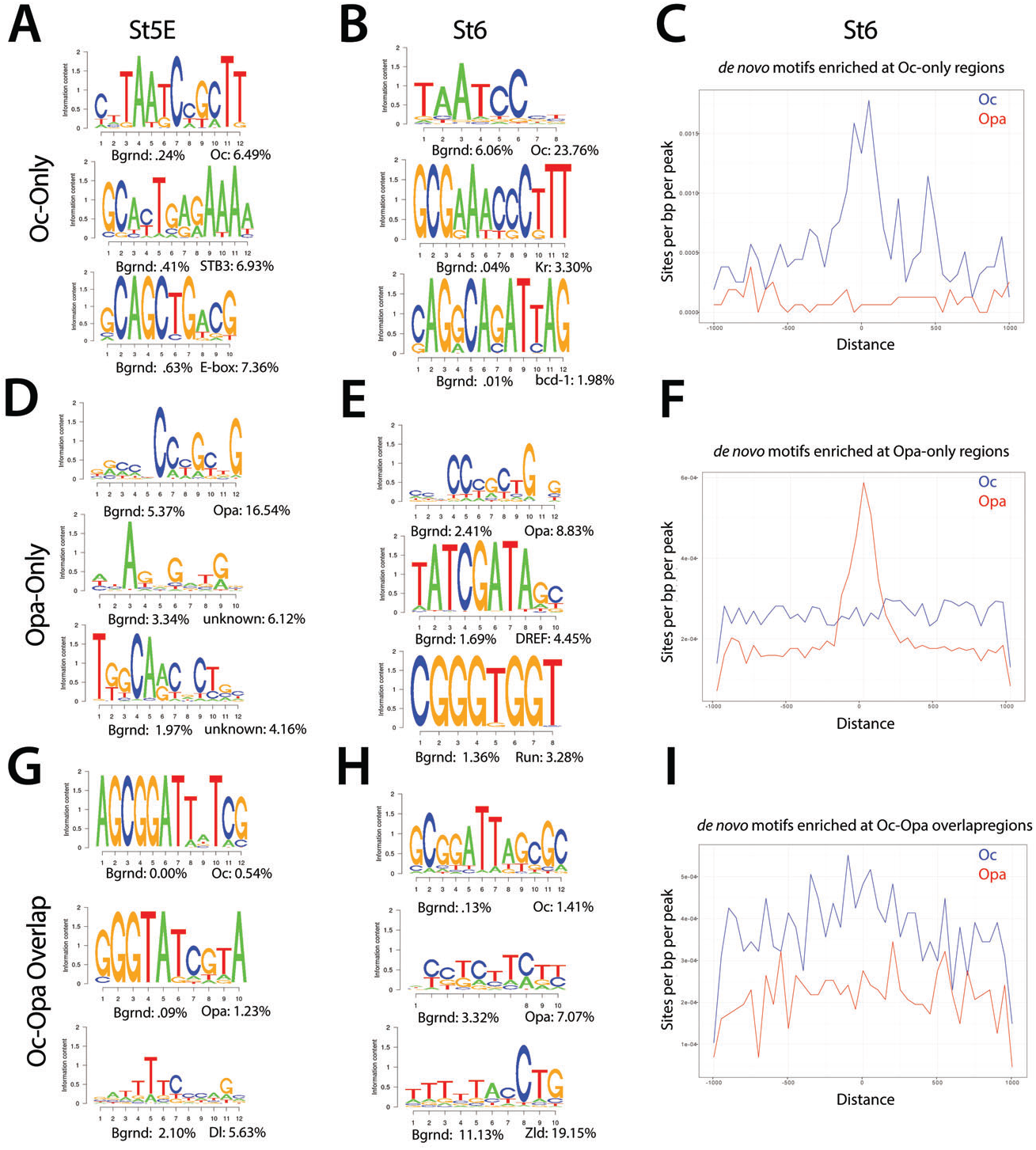
Enrichment of Opa and Oc *de novo* motifs in subsets of peaks that correspond to Opa-only, Oc-only, or Opa/Oc-bound regions identified by ChIPseq. (A,D,G) Stage 5E HOMER de novo motif analyses for Oc only (no Opa) (A), Opa only (no Oc) (D), and Opa and Oc only (G) peaks. (B,E,H) Stage 6 HOMER de novo motif analyses for Oc only (no Opa) (B), Opa only (no Oc) (E), and Opa and Oc only (H) peaks. (C,F,I) Enrichment plots of Stage 6 Oc only (C), Opa only (F), and Oc-Opa overlapping peak (I) motifs at Oc (blue) or Opa (red) peaks.

To further investigate Oc and Opa binding dynamics we compared the same predicted motifs against only those peaks from the previous analyses which did not overlap with Zld peaks. Removing Zld overlapping peaks produced negligible changes early, but resulted in a marked coalescence of Oc-only motifs around the remaining Oc peaks (Supplemental Figure 2.2A-F) demonstrating lower Oc motif density at Zld-binding loci at the late stage (St6).

A feed forward relay from Bcd to Oc has previously been shown, so we performed *de novo* motif analysis on publicly available Bcd ChIPseq data to compare to Oc motif analysis at St5E when the two factors are most co-expressed (Supplemental Figure 2.1A and Supplemental Figure 2.2G). Unsurprisingly, the predicted motifs are highly similar, further supporting the published concept of a Bcd-to-Oc handoff. We further performed *de novo* motif analysis of those Oc peaks which do not overlap with Bcd peaks and did not find any major differences to the Oc-only motif analysis (data not shown).

### Opa and Oc overlaps in a narrow temporal window during embryogenesis

Having determined that *opa* and *oc* expression domains overlap, we sought to confirm that this overlap results in expression of both transcription factors within individual cells in the overlapping region. Using super resolution microscopy we were able to image individual allele transcription of both *oc* and *opa* within individual nuclei of the overlap region (Figure 3A). We were further able to confirm that the overlapping region diminishes as cellularization ends and gastrulation begins by counting cells along the AP axis coexpressing both transcripts (Figure 3B). This finding is intriguing as this overlapping domain resides in the region of the embryo which will eventually beget the nascent brain and implies the potential for dual binding of these two activators uniquely within these cells.

**Figure 3.**
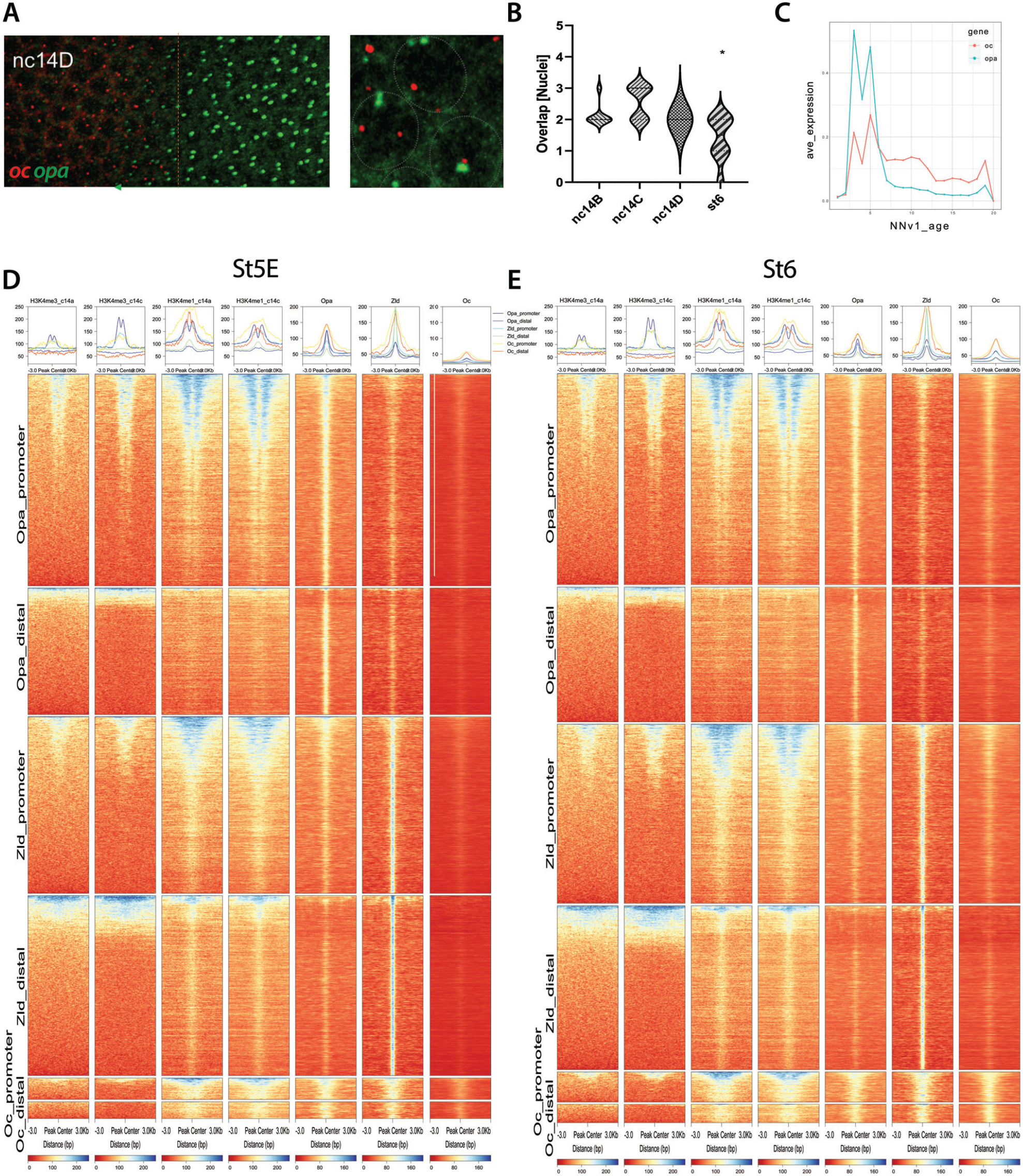
Opa/Oc subcellular protein and transcriptional dynamics. (A) Super resolution microscopy reveals simultaneous transcription of *opa* (green arrow region) and *oc* (bounded by orange dotted line) within individual nuclei (grey dotted lines) in the *opa/oc* overlap region. (B) Nuclei counts of *opa/oc* overlap region width by stage from the onset of *opa* expression to St6. The overlapping region is significantly lower at St6 than any of the cellularization stages (p<.05 compared to all other individual stages). (C) Neural network age prediction plot of *oc* and *opa* expression from publicly available scRNAseq datasets^24,29^. X-axis values are neural network age prediction pseudotimes between ages 0hr and 20hr. (D,E) Stage 5E (D) and Stage 6 (E) Opa, Oc, and Zld ChIPseq peak correlation by promoter or distal enhancer subclusters to common histone marker, Opa, Oc, or Zld ChIPseq peak loci.

To better characterize the temporal dynamics of *opa/oc* expression overlap, we analyzed publicly available single cell RNA sequencing (scRNAseq) data spanning 1-7hr into embryonic development (∼St3-11) (Supplemental Figure 3.1B-E). We further used neural network age prediction of the transcriptomic temporal landscape^24^ to visualize *opa/oc* expression (Supplemental Figure 3.1F). In support of our findings that the *opa/oc* overlapping region is transient, we found that *opa* and *oc* expression peaks at approximately the cellularization/gastrulation transition (Figure 3C). Intriguingly, we also found that *opa* and *oc* expression drops significantly early in gastrulation as well with *oc* diminishing much more gradually than *opa* and a small population (45 cells) of *opa/oc* coexpressing cells between 1-3 post fertilization (Figure 3C, data not shown).

To examine the potential for Opa and Oc cooperation during this stage of development, we compared St6 chromatin occupancy of Opa and Oc between gene loci based on their published expression shifts following *opa* knockdown^3,23^. Oc only peaks reside predominantly at gene loci insignificantly changed by *opa* KD as do Opa peaks (Supplemental Figure 3F). However, Oc peaks broadly correlate most strongly with gene loci of genes which increase in expression following *opa* KD contrary to Opa only peaks which associate most strongly with insignificantly affected gene loci (Supplemental Figure 3F,G). Intriguingly, together, this suggests a small number of genes in some cells coexpressing Opa and Oc may be conversely regulated by these two transcriptional activators.

We next sought to investigate potential differences in Opa and Oc peak genomic distributions. Both transcription factors correlate with the transcriptional silence marker, H3K4me3, early and late peak locations similarly between St5E and St6 at gene promoters but not distal enhancers (Figure 3D,E, Supplemental Figure 3C-H). Similarly, both transcription factors bind similarly between stages 5E and 6 to genomic loci marked by H3K4me1 regardless of whether the loci were marked prior to, after, or during the peak binding (Figure 3D,E). Curiously, at promoter loci, Oc and Opa appear to occupy complimentary niches relative to these histone marks, with Opa binding coalescing bimodally around the histone marks and Oc peaks centering atop them, reflecting a more Zld-like profile for Oc than Opa (Figure 3D,E, Supplemental Figure 3C-H). Strikingly, Oc promoter and distal enhancer peaks increase dramatically between St5E and St6 at Zld peak loci (Figure 3D,E, Supplemental Figure 3.2).

### Opa and Oc overlap is likely involved in spatiotemporal localization of downstream AP and DV gene expression in that region

Further, to interrogate the regulatory dynamics of Opa and Oc, we looked at peak overlap between Opa, Oc, Bcd, and Zld. Nearly all Oc peaks overlap with at least Opa or Zld at both St5E and St6 (89% early, 96% late; Figure 4A). Interestingly, there is a large difference between distal enhancer and promoter loci in this regard wherein Oc only peaks are more than twice as frequent in distal enhancers than promoters (Suplemental Figure 4K-N). Additionally, this reduction in Oc only peaks at promoters seems to be nearly entirely driven by overlap, especially St6 overlap, with Zld peaks. Together these data suggest that Oc may play a more independent role at distal enhancers than promoters where Oc, similar to Opa^3,23^, may be acting as a regulatory substitute for waning Bcd and Zld factors.

**Figure 4.**
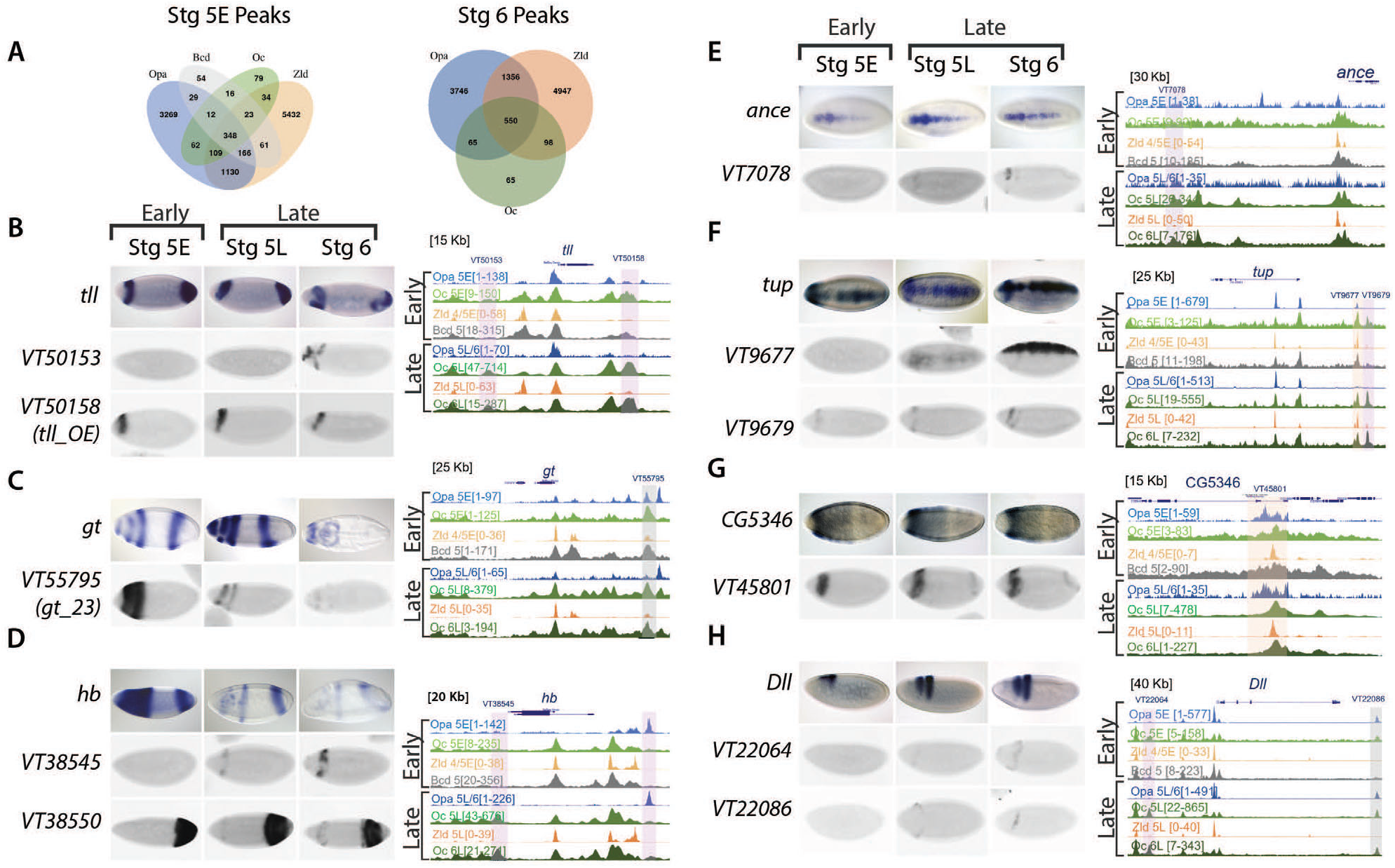
Oc ChIPseq data demonstrates binding in both AP and DV axis, including some stage 5 enhancers, as well as other later acting enhancers. (A) Venn diagrams of Opa, Oc, Bcd, and Zld peaks at distal enhancers at stages 5E and 6. (B-H) ChIPseq data and enhancer expression patterns for important developmental factors. ChIPseq datasets: Opa at stage 5 early (light blue), Oc at stage 5 early (light green), Zld at stage 4/5 early (light orange), Bcd at stage 5 (grey), Opa at stage 5 late/ stage 6 (dark blue), Oc at stage 5 late (olive green), Zld at stage 5 late (dark orange), and Oc at stage 6 late (dark green) from previous studies were aligned using UCSC genome browser. Numbers in square brackets indicate maximal peak heights and colored highlights marking the peaks indicate enhancers of interest with different occupancy in our study. The grey highlights mark Oc/Opa binding, light purple indicates Opa or Oc individually bound regions, and orange highlights mark Zld-bound enhancers. For each panel (B-H), endogenous expression patterns for (B) *tailless* (*tll*), (C) *giant* (*gt*), (D) *hunchback* (*hb*), (E) *angiotensin converting enzyme* (*ance*), (F) *tailup* (*tup*), (G) *CG5346*, and (H) *distal-less* (*Dll*) are extracted from publicly available Fruitfly database, and expression patterns of highlighted enhancers (Vienna tile ID in blue above).

We next explored enhancer occupation by Opa, Oc, Bcd, and Zld at targets with distinct expression ear the Opa/Oc overlap domain or known to play major roles in brain and neuroblast development in Drosophila melanogaster. Oc and/or Opa occupy enhancers near giant (gt), tailless (tll), hunchback (hb), distal-less (Dll), Angiotensin converting enzyme (ance), tailup (tup), eyeless (ey), CG5346, empty spiracles (ems), buttonhead (btd), toy, H6-like-homeobox (hmx), disheveled (dsh), kdm5, eyeless (ey), ventral nervous system defective (vnd), posterior sex combs (psc), and acheate (ac) (Figure 4B-H & Supplemental Figure 4A-J). Those Oc and/or Opa occupied enhacers at these genes with archived gene trap expression patterns in the StarkLab database^32^ reveal a clear pattern of expression at or near the Opa/Oc overlap domain (Figure 4B-H). Intriguingly, several genes being regulated in the overlap region are expressed in a DV pattern suggesting this Opa/Oc co-regulation is not limited to AP patterning (Figure 4E-H).

To experimentally test these analyses of Opa and/or Oc target genes, we sought to experimentally reproduce a result from our analyses. To this end, we chose to epigenetically modulate Oc-mediated *hb* expression. Using *oc-RNAi*^33^ to knock down Oc levels, we were able to eliminate, at St5L when Opa and Oc are no longer simultaneously available to potentially compensate for one another, the second band, located in the region of *opa/oc* co-expression, of anterior *hb* expression which can be seen exogenously reproduced via enhancer driven *lacZ* from the StarkLab database (see Figure 4D, Figure 5B,C, Supplemental Figure 5).Together, these data suggest dynamic roles for Opa and Oc in gene regulation which include a clear potential for establishment of a head lineage niche beginning within their early, transient overlapping region.

**Figure 5.**
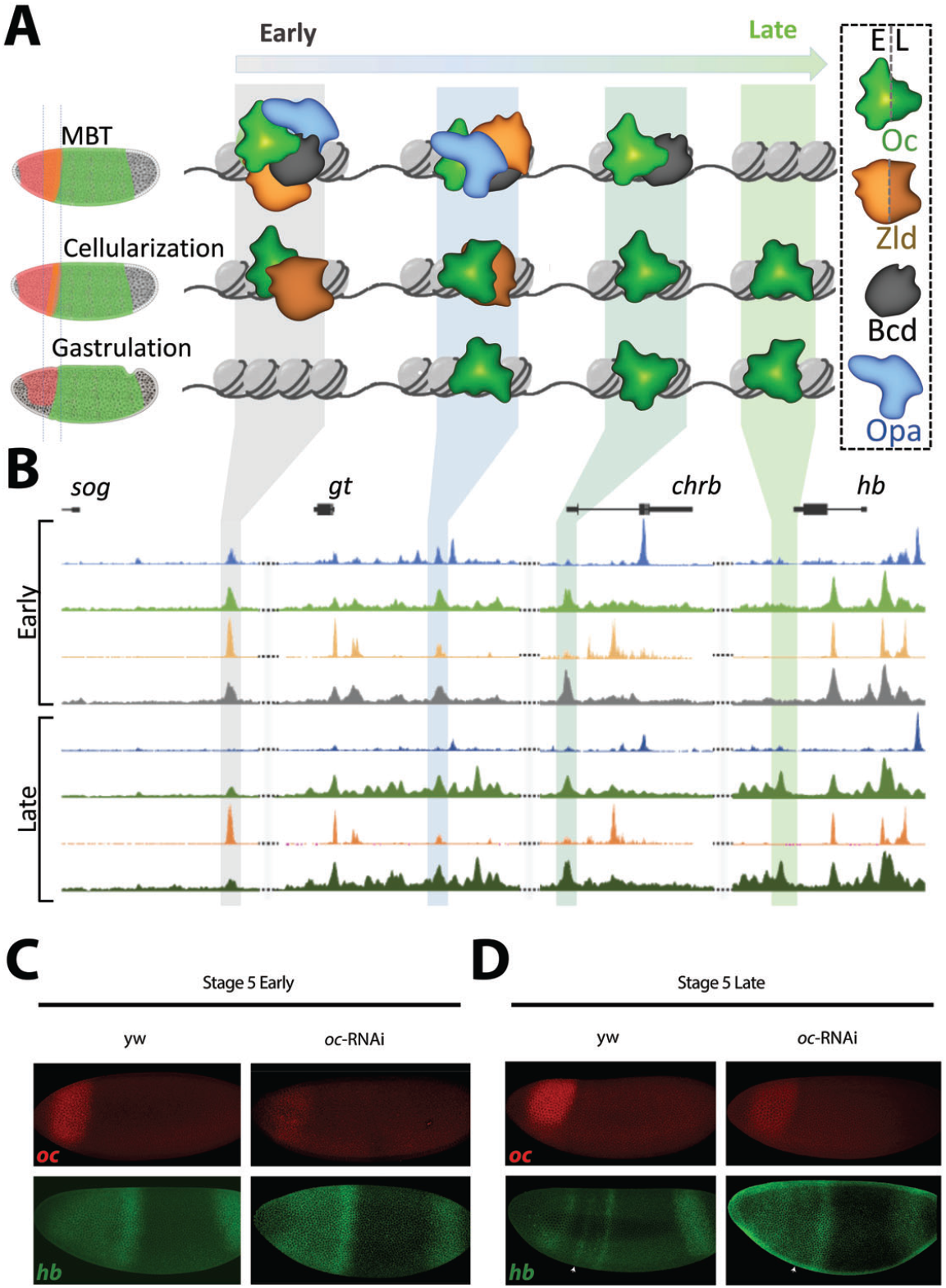
Oc and Opa play diverse roles in embryonic head development. (A) Illustration of proposed Opa/Oc action during MZT. Leading into cellularization, Opa and Oc tend to share enhancer occupancy with maternal transcription factors, e.g. Zld, Bcd, as cellularization progresses, Opa and Oc switch to less cooperative binding with maternal factors, and ultimately shift to distinct roles as gastrulation begins and *opa/oc* expression domains divorce. (B) ChIPseq occupancy plots of example regions of significant relevancy in head region development. (C-D) *oc* depletion by shRNAi results in an absence of the second *hb* band (white arrowheads) late in cellularization when Oc ChIPseq shows Oc occupancy, complimenting the *V38545* enhancer expression pattern shown in Figure 4D.

## Discussion

In this study, we combine meta analyses and *in vivo* experimentation to reveal Opa/Oc epigenetic dynamics during early embryonic development. We demonstrate an overlap in Oc and Opa expression in cells likely destined for brain development before the onset of gastrulation and lasting less than an hour. During this short period of overlap, we noticed that both Opa and Oc binding sites on the genome are less resolved than after their expression domains suggesting the possibility of competitive or cooperative binding between the two factors. This implication was further supported by our finding that a broad majority of Opa and Oc peaks overlap on the genome. Together, these early findings strongly point toward a developmental instrument for the specification of a cell niche concurrent with or preceding the onset of gastrulation.

Opa and Oc binding motifs identified via Homer *de novo* motif analysis of mutually exclusive peaks from publicly available ChIPseq datasets demonstrating individual temporally shifting binding dynamics of Opa and Oc. Interestingly, we found association of both Opa and Oc peaks with chromatin architecture factors, Trl/Gaf and Dref-1/Beaf-32, binding motifs^34^. However, inspection of publicly available ChIPseq data^35^ revealed that Gaf does not appear to bind at Opa/Oc regulated enhancers for genes investigated in this study (data not shown). Further investigation is needed to untangle the intriguing implications of this finding and to determine whether Opa or Oc peaks are involved in TAD insulator functions. Through investigation of ChIPseq datasets, we were further able to identify relative shifts in Opa and Oc binding to Zld peaks at promoters rather than distal enhancers, supporting a model whereby Opa and Oc regulate transcription at late enhancer regions as pioneer-like factors (Supplemental Figure 4K-N). However, whether sequential binding of the two proteins to the same DNA motif is related to cell specification, and whether Oc activates a third wave of zygotic transcription in brain cells needs further investigation.

Expression of some genes is unchanged when Opa protein is diminished^3^ despite broad and abundant genomic occupancy by Opa, implying cooperative and compensatory transcriptional regulation with other transcription factors of similar regional spatiotemporal abundance, such as Oc. The expression pattern of both Opa and Oc is dynamic at stage 5, but we characterize here, using whole embryo and super resolution microscopy coupled with both manual and automated quantification techniques, as well as scRNAseq analysis, the transient Oc/Opa overlap suggested early in our study. Much of the epigenetic landscape remains unexplained at the cellularization/gastrulation transition and these Opa/Oc dynamics are undoubtedly involved; future studies are needed to investigate how the embryo leverages this unique cell niche to pattern the brain/head.

We further investigated how Opa and Oc are associated and regulate the zygotic genome in both Anterior-Posterior (AP) and Dorsal-Ventral (DV) axes at cellularization and at gastrulation. We found that these associations tend to correspond with binding at enhancer regions which drive expression in bands at or near the Opa/Oc overlapping domain. Rather than transcriptional activation being linked directly to absolute Oc concentration, Opa may act to modulate Oc’s effective concentration: e.g. lower levels of Oc may be required to activate enhancers bound by Opa. This postulation would explain our findings that some Oc-only peaks are capable of driving narrow band expression in this region rather than across the entire *oc* expression domain and why knockdown of *oc* was sufficient to eliminate *hb* expression there as well. However, future work is needed to determine if this phenomenon is driven by Opa regulation of Oc levels, whether Opa plays a compensatory or cooperative role at Oc peaks, or some mixture of these and whether these regulatory dynamics are direct or indirect.

In summary, we observed Opa/Oc expression and binding dynamics through cellularization and up to early gastrulation characterized by Opa/Oc transient overlap and suggestive of a mechanism whereby Opa and Oc appropriate epigenetic control of some genes from maternal factors early, work together transiently, and diverge in regulatory function with Oc outlasting Opa expression and driving new expression late (Fig. 5A).

## Methods

### Fly stocks and husbandry

Wild type flies used in this study were of the yw [67c23] strain. Flies were reared under normal conditions at 23°C, with the exception of UAS-RNAi lines crossed to Gal4^36^ and yw control flies for those experiments which were incubated at 26.5°C. Virgin ocRNAi (Bloomington Drosophila Stock Center (BDGP) #34327, #29342) females were crossed to matzyg.Gal4 or MTD.Gal4 males (ref or BDSC#, #31777). Depletion of oc was achieved by crossing the virgin females from this cross to ocRNAi males.

### In situ hybridization, imaging, and analysis

Standard protocols were used for 2-4hr embryo collection, fixation, and staining. FISH was performed using antisense RNA probes labeled with digoxigenin-, biotin-, or DNP-UTP to detect transcription of target genes.

Images were acquired using a Zeiss LSM 900 confocal and super resolution microscope. Confocal images were taken using a 20x air lens and super resolution, Airyscan, images were taken using a 40x water objective using 488nm, 561nm, and 647nm lasers.

Image processing was performed in Fiji (ImageJ) using standard z-projection procedures. Processed images used for expression domain profiling were then used to create segmentation masks in Ilastik^16,37^. To generate the expression domain plots, a Python script was used to average pixel fluorescent intensity for each channel in 10px-wide slices along the AP axis of the embryos; these data were then plotted in GraphPad Prism.

### Bioinformatics

Oc and Bcd ChIPseq bed peak files of dm3 coordinates were converted to dm6 using UCSC liftOver tool. Oc and Bcd ChIPseq bigwig signal traces were converted from dm3 to dm6 assembly using crossMap (v0.6.4, PMID: 24351709). Opa and Zld processed data (dm6 assembly) were collected from our previous study (PMID 32701060).

To understand overlapping of different transcription factor binding sites across the genome, peak regions were combined and overlapping peaks were merged. Combined regions that overlapped both Opa and Oc peaks were defined as Opa-Oc overlap regions; regions overlapping with either Opa or Oc peaks were defined as Opa-only and Oc-only regions respectively. Region overlap analysis was performed using bedtools (v2.30.0) and Venn diagrams were generated using VennDiagram R package. Further de novo motif analysis was performed on different ChIPseq regions using the HOMER program (PMID 20513432) with default parameters and with options -size 200 and -mask. Selected de novo motifs identified from peak regions were queried against the Opa-Oc overlap, Opa-only and Oc-only regions for comparison and for generating motif aggregation plots, with the -size 2000 -hist 50 options. DNA sequence logos were plotted using the seqLogo R package. ChIPseq peak regions were associated with nearest gene transcription start sites using the annotatePeaks.pl module of HOMER. Promoter peaks and distal peaks were distinguished using a distance cutoff of 3kb to the nearest transcription start sites.

In addition, computeMatrix and plotHeatmap modules of deepTools (v3.2.1) were used to calculate and plot normalized histone mark and transcription factor signal intensities surrounding selected ChIPseq regions. For this and all subsequent data presented using heatmaps, the first sample in the heatmap was used for sorting the genomic regions based on descending order of mean signal value per region; all other comparison samples were plotted using the same order determined by the first sample. IGV browser (PMID 21221095) was used to visualize ChIPseq signals at individual loci.

ChIPseq peak-associated genes and RNAseq differentially expressed genes were subjected to overlapping statistical analysis (Fisher’s exact test), and the results were presented in overlap gene count and overlap p-value heatmaps.

Publicly available single-cell RNAseq data was downloaded from GEO database (GSE190147). The processed gene count table of a total of 547,805 single nuclei from stages drosophila embryos was subject to downstream analysis. As note, each single nucleus was assigned with a developmental age score (NNv1_age) using neural network-based prediction^24^.

To track gene expression across developmental stages, age score (NNv1_age) of each nucleus was rounded up to the nearest integer to calculate average gene count values for the nuclei at the same developmental stage (NNv1_age_bin). Gene expression values across 20 different time points were presented in a line plot.

To explore co-expression of two genes at single cell level, double positive nuclei (at least 1 count for both genes) were separated from other nuclei. A violin plot was presented to show cell distribution across developmental age for both positive and negative groups.

Unless noted otherwise, R was used to calculate statistics and generate plots.

### ChIPseq procedure and analysis

ChIPseq was used to determine the binding sites of transcription factors and other chromatin-associated protein in the genome and to understand how proteins interact with the genome to regulate the gene expression in Drosophila embryo. ChIPseq libraries were generated from the University of California, Santa Cruz (UCSC) genome browser platform. The ChIPseq reads from previous studies were aligned were aligned to Drosophila reference genome assembly (UCSC dm3) (Datta et al. 2018) at different timepoints: Opa at stages 5 early and 5 late/stage 6, Oc at stages 5 early, 5 late, and 6 late, Zld at stages 4/5 early and 5 late, and Bcd at stage 5. The resulting alignment tracks helped us to detect important genomic regions to study the mentioned factors.

## Acknowledgements

We are grateful to Rhea Datta and Angela Stathopoulos for generously providing us with fly lines, antibodies, and ChlPseq data. We would further like to thank Anupama Chandrasekhar for her help with the StarkLab database; Hinduja Sathishkumar and Saubia Zareen, students in the Koromila Lab, for their assistance with administrative tasks and fly husbandry; and Mounia Lagha for helpful discussions. This work was made possible by funding from the UTA STARS program and the Bioinformatics Resource Center at the Beckman lnstitute of Caltech.

## Data and Code Availability

The code and pipelines generated during this study is available at GitHub (https://github.com/gaofan83/Koromila_group).

## Notes

### Competing Interest Statement

The authors have declared no competing interest.

